# Respiratory syncytial virus activates Rab5a to suppress IRF1-dependent IFN-λ production, subverting the antiviral defense of airway epithelial cells

**DOI:** 10.1101/547182

**Authors:** Leiqiong Gao, Wei Tang, Jun Xie, Sisi Chen, Luo Ren, Na Zang, Xiaohong Xie, Yu Deng, Enmei Liu

**Author notes:** **Corresponding author**: Enmei Liu, MD, PhD, Department of Respiratory Medicine Children’s Hospital of Chongqing Medical University, Chongqing, China, 400014.

## Abstract

Human respiratory syncytial virus (RSV) is a negative-strand RNA virus that causes severe acute pediatric respiratory tract infections worldwide. The limited effective antiviral options and lack of an effective vaccine against RSV highlight the need for a novel anti-viral therapy. One alternative is to identify and target the host factors required for viral infection. All viruses, including RSV, utilize cellular trafficking machinery to fulfill their life cycle in the infected host cells. Rab proteins mediate specific steps in intracellular membrane trafficking through the recruitment and tethering of fusion factors, and docking with actin- or microtubule-based motor proteins. Using RNA interference to knock down Rab proteins, we document that the micropinocytosis-associated Rab5a is required for RSV infection. RSV infection itself induces activation of Rab5a, and inhibition of this activation reduces RSV infection, but the mechanism for this effect remains unknown. Interferon (IFN) signaling plays an important role in innate immunity, and recent studies have identified IFN-λ (lambda), a type III IFN, as the most important IFN for antiviral immune in response to RSV infection of mucosal epithelium. However, how the RSV-induced Rab5a suppresses airway epithelial antiviral immunity has not been unraveled. Here, we show that activated Rab5a inhibits IRF1-induced IFN-λ production and IFN-λ-mediated signal transduction via JAK-STAT1, thereby increasing viral replication. Rab5a knockdown by siRNA resulted in stimulation of IRF1, IFN-λ and JAK-STAT1 expression, and suppressed viral growth. Our results highlight new role for Rab5a in RSV infection, such that its depletion inhibits RSV infection by stimulating the endogenous respiratory epithelial antiviral immunity, which suggests that Rab5a is a potential target for novel therapeutics against RSV infection.

**Author summary:** RSV is the leading cause of lower respiratory tract infection in under 5 years old children. Worldwide. We identified Rab5a as a host factor involved in RSV infection via RNA interference to knock down familiar Rab proteins in human lung epithelial A549 cells infected with RSV. Rab5a belongs to Rab GTPases subfamily, which contributes to intracellular trafficking to promote virus infection. Knockdown or inactive (GDP-bound) Rab5a results in low infection and replication through stimulating IRF1, IFN-λ and JAK-STAT1 expression, and suppressed viral growth. Besides, we propose that the regulation of Rab5a expression during RSV infection might be a viral strategy to promote its infectivity.

## Introduction

Human respiratory syncytial virus (RSV) belongs to the *Paramyxoviridae* family [1], and is a leading cause of respiratory tract infection in young children [2]. Approximately 4 million children worldwide are admitted to hospitals each year with RSV infection, 3.4 million of whom develop severe symptoms such as bronchiolitis and pneumonia [3–5]. The healthcare costs of hospitalization from the RSV-infected patients are significant [6,7], and despite years of ongoing efforts, there is currently no safe or effective vaccine available to protect children and minimize the global burden of RSV. Thus, identification of host factors required for RSV infection may be considered as a plausible alternative to develop a therapeutic regimen.

Being obligatory intracellular parasites, viruses utilize diverse cellular trafficking machinery to achieve productive life cycles in the infected host cells. Members of the Rab family of cellular proteins regulate actin- or microtubule-based motor proteins and intracellular membrane trafficking, and have been implicated in various steps of the viral life cycle, including replication, assembly, and budding. To identify cellular Rab proteins required for RSV infection, we interrogated the role of nine widely expressed Rab proteins (Rab1a, Rab2a, Rab4a, Rab5a, Rab6a, Rab7a, Rab8a, Rab9a, Rab11a) that are involved in the endo- or exocytic pathways. Using specific siRNA to knock down each Rab protein, we found that the micropinocytosis-associated Rab5a protein is required for RSV infection, which confirmed and extended our previous iTRAQ data suggesting this role of Rab5a (unpublished data). Additionally, RSV infection activated Rab5a, which is related to actin-mediated micropinocytosis of RSV, and recent studies showed that inhibition of Rab5a results in decreased RSV titers [8]. In another study, depletion of Rab5a by siRNA resulted in decreased RSV replication, but viral binding was not affected [9]. Together, these findings further suggested that inhibition of Rab5a activation affects post-entry steps of the viral life cycle.

Rab5a, a major member of the small GTPase Rab family, is mainly localized to the cytosolic face of the plasma membrane, early endosomes, clathrin-coated vesicles and macropinosomes [10–13]. The Rab5a activity depends on GDP/GTP association [14]; the activity is also spatially regulated and ensure the bi-directionality of the processes it governs. Several studies have demonstrated that Rab5a, in particular, plays a critical role in viral infection. For example, components of positive-strand RNA viral replication can hijack Rab5a to promote viral replication [15]; the influenza A virus uses Rab5a to modulate Annexin-A1, thus enhancing its replication [16]; HIV and hepatitis B/C viruses co-opt Rab5a to enter cells [17–21]. The involvement of Rab5a in RSV endocytosis or micropinocytosis has been described previously [8], and as mentioned above, knockdown of Rab5a resulted in decreased RSV replication [9]. In parallel, several studies demonstrated that Rab5a is closely related to innate immunity. IFN-gamma (IFN-γ), a type II interferon, induces Rab5a synthesis [22–24], which acts to mediate the bactericidal effect of IFN-γ. In addition, IL-4 and −6 increase Rab5 expression [25], and IL-4 also extends the retention of Rab5 on phagosomes in a PI3K-dependent manner [26]. Rab5a is closely related to the IFN-signaling JAK-STAT pathway, and downregulation of Rab5a increases STAT1 expression [27,28]. Rab5a also affects TLR4-mediated innate immunity [29]. Rab5a is required for the formation of the early endosome, which is related to the IFN-induced transmembrane proteins of the IFITM family; moreover, the type I IFN receptor complex is also differentially sorted at the early endosome [30]. Taken together, these studies suggest that Rab5a may affect the innate immunity in RSV infection. Lastly, several RNA viral nonstructural proteins, such as the NS proteins of RSV, subvert IFNs, and Rab5 has been shown to co-localize with NS-induced structures of the SFTS (Severe fever with thrombocytopenia syndrome) virus [31]. Based on these findings, we hypothesize that Rab5a facilitates RSV infection, not because it promotes virion binding, but because it inhibits the cell-intrinsic antiviral IFN pathway. Nonetheless, there are currently no studies of the effect of Rab5a on IFN signaling.

As mentioned, IFN signaling is a major arm of the innate antiviral response of the host. Recent studies revealed that that IFN-λ, a type III IFN, is also an important IFN of the airway epithelium [32,33]. Further studies suggested that type I IFNs (i.e., IFN-α and IFN-β) are critical for the clearance of infection, whereas IFN-λ is the most important IFN regulating mucosal epithelial cell responses to viral infection [33,34]. Recent studies of the Koff group found that IFN-λ is the first produced IFN of the RSV-exposed nasal epithelium [35]. Moreover, RSV could inhibit IFN-λ production in lung epithelial cells, and IFN-λ was critical for antiviral immunity to RSV [36,37]. Further studies suggested that RSV induces IFN-λ production by activating IRF1, a transcription factor for the IFN-λ gene [35]. However, the potential role for Rab5a pathway in modulating IFN-λ and its related innate immunity in RSV infection has not been reported. Here, we have explored the effect of the Rab5a pathway on RSV infection in airway epithelial cells and the role of IFN-λ-related factors in this process. We show that RSV infection indeed activates Rab5a, which in turn facilitates viral infection by regulating the pathways related to IFN-λ.

## Results

### Rab5a is essential for RSV production

Studies in this section interrogated the effect of Rab5a on RSV infection in airway epithelial A549 cells. To this end, we depleted Rab1a, Rab2a, Rab4a, Rab5a, Rab6a, Rab7a, Rab8a, Rab9a and Rab11a by treating the cells with specific siRNAs (or control siRNA) for 24 h p.i., followed by infection with purified RSV A2 and incubation for another 36 h. Depletion of Rab proteins was confirmed by immunoblotting analysis of the total cell lysates (Figure 1B). To determine the viral load, the viral N gene RNA was quantified by RT-qPCR on RNA isolated at 36 h p.i. (Figure 1C). The syncytial area and number were counted using Image J (Figure 1A & 1D). Little alteration of RSV replication was observed following depletion of Rab1a, Rab2a and Rab6a compared to the control cells treated with siRNA (Figures 1A and 1C). A moderate decrease in RSV replication was observed on knockdown of Rab4a, Rab7a, Rab8a and Rab9a, with 40-70% less N gene detected both in the cells and supernatant compared to the control. The effects of Rab11a depletion were similar to those reported previously [38,39]. The most potent effect on RSV propagation was observed in cells depleted of Rab5a. Indeed, after 48 h p.i., only a few infected cells were observed in Rab5a-depleted cultures (Figure 1A & 1D) and the level of N was more than 30-fold lower than the control (Figure 1C). These results suggest that Rab5a plays an important role in RSV propagation.

**FIGURE 1.**
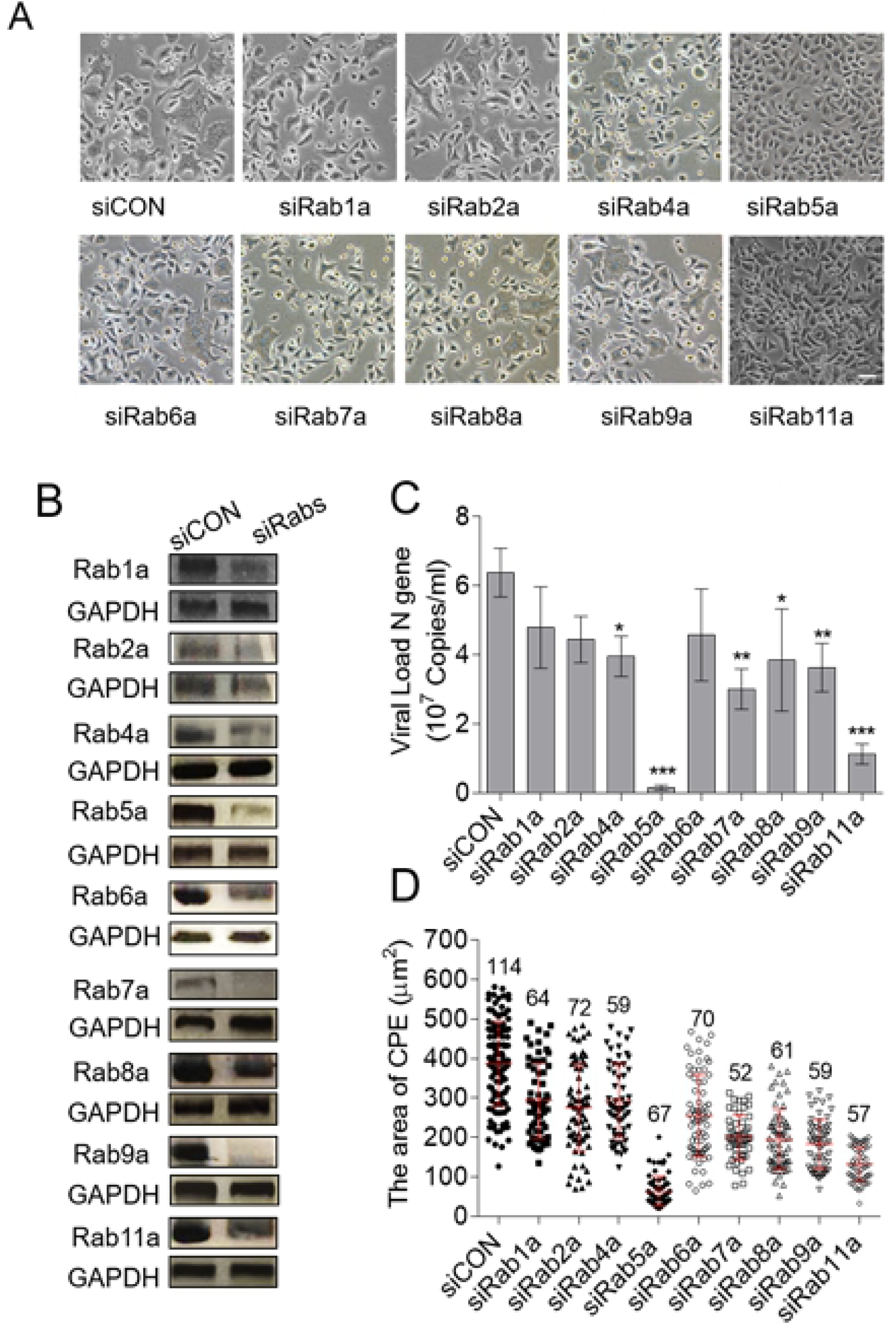
Rab5a depletion reduces RSV propagation. A549 cells were transfected with either siRNA control (siCon) or specific siRNAs targeting Rab proteins (siRabs), and then were infected with RSV as described in Materials and Methods. (A) Bright field image of infected cell cultures, taken at 36 h p.i. Bar = 20 μm. (B) Analysis of efficiency of Rab protein depletion. (C) RSV propagation was scored by measuring the amount of RSV both in attached cells and released into the culture supernatants by RT-qPCR. (D) The graph shows the comparative average size of RSV syncytia with siRab1a (n=64), siRab2a (n=72), siRa4a (n=59), siRab5a (n=67), siRab6a (n=70), siRab7a (n=52), siRab8a (n=61), siRab9a (n=59), siRab11a (n=57), relative to that siCON (n=114) from 20 different fields using ImageJ. Bars represent the means±SD for three independent experiments, each performed in duplicate. P values were calculated based on unpaired Student’s t-test between siCON and siRabs. Significant results (***, *p*<0.001, **, *p*<0.01 and *, *p*<0.05) are indicated.

### RSV infection increases Rab5a expression

To investigate the induction of Rab5a by RSV in the infected cells, we measured total Rab5a mRNA and protein by RT-qPCR and immunoblotting at various post-infection times. It was observed that both Rab5a mRNA and Rab5a protein levels significantly increased starting at 1 h p.i. (Fig. 1), when compared with mock infection. These results confirmed that RSV indeed induces Rab5a expression.

### Rab5a-GTP active form required for RSV infection

To examine whether Rab5a-GTP affects RSV replication, recombinant EGFP-Rab5a (wild type), constitutively active (C/A) mutant, Q79L, and the dominant negative (D/N) mutant, S34N, were transiently transfected in A549 cells. Q79L can enlarge Rab5a-positive vacuoles, but fails to undergo further maturation[40]. S34N inhibits Rab5a-GDP transfer to Rab5a-GTP, and thus inhibits Rab5a located on the membrane[41]. After 24 h transfection, cells were infected with RSV for the indicated time. Viral load analysis demonstrated that S34N was the only one that caused a significant decrease in RSV infection when overexpressed (Fig. 3B). These results confirm and extend those reported by the Helenius laboratory [8]. Consistent with the viral load results, we also found that Q79L showed higher co-localization with RSV compared to EGFP-Rab5a cells by using confocal microscopy. In contrast, EGFP-Rab5a S34N showed the lowest amount of virus particles and least co-localization among the three groups (Fig. 3A). However, no significant changes were observed in cells expressing EGFP-Rab5a. Together, these results confirmed that the active form of Rab5a is in fact essential for RSV replication in vitro.

**FIGURE 2.**
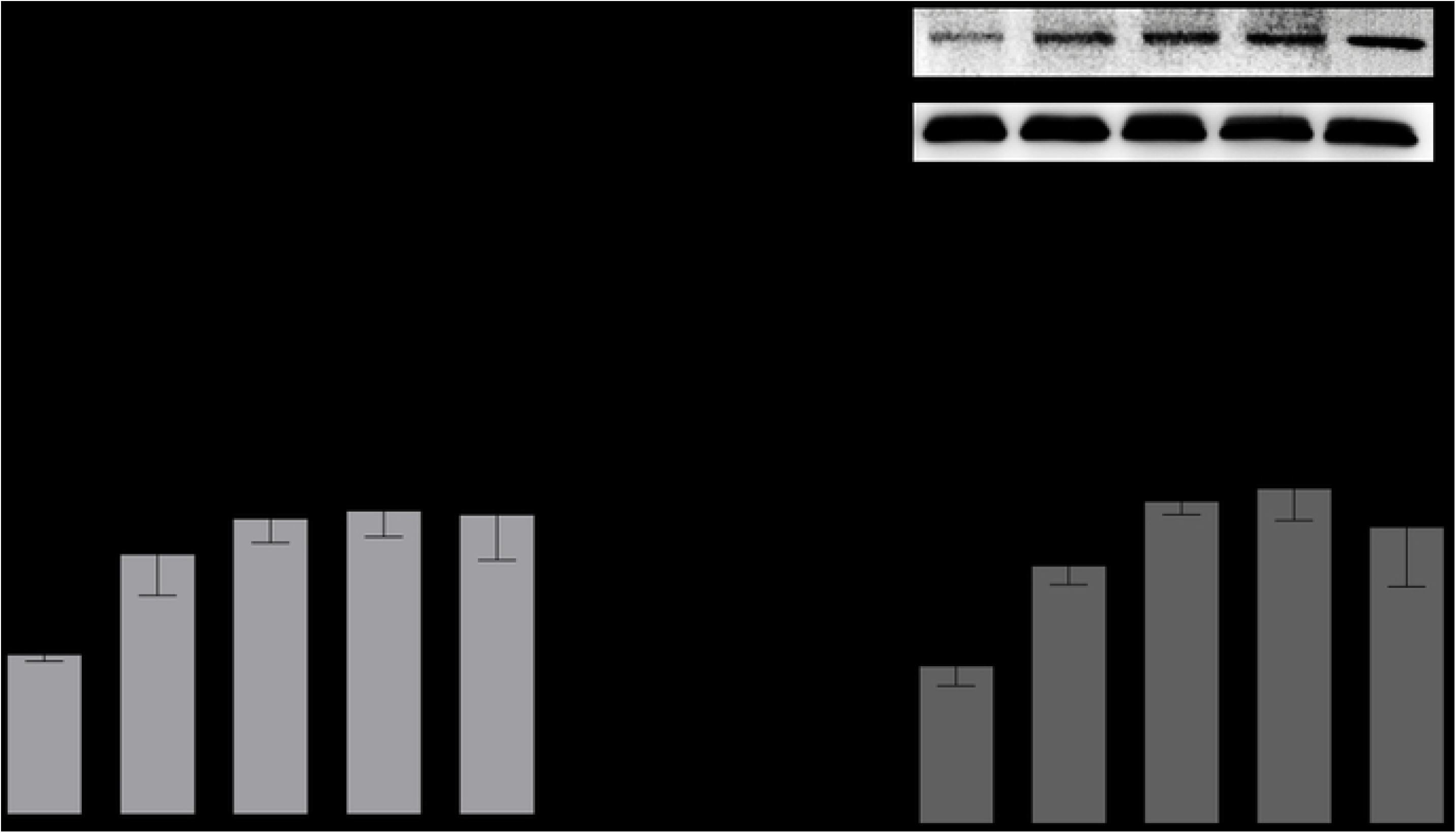
RSV increases Rab5a expression at early infection times. Nasopharyngeal aspirates (NPA) from 26 RSV-infected infants and 24 uninfected controls were used and processed as described in Materials and Methods. Total protein was extracted from the cell pellets. (A) Rab5a protein expression in NPAs. A549 cells were infected with RSV and at indicated times (0, 1, 2, 6 12 h p.i.), samples were collected for measurement of Rab5a mRNA and protein expression. (B) Rab5a mRNA and (C) protein expression at various post-infection times. (D) Semi-quantitative analysis of the data from (C), using Image J. Bars represent mean ± SD for three independent experiments performed in duplicate. P values were calculated based on unpaired Student’s t-test between Control and RSV infection. Significant results (***, *p*<0.001, **, *p*<0.01 and *, *p*<0.05) are indicated.

**FIGURE 3.**
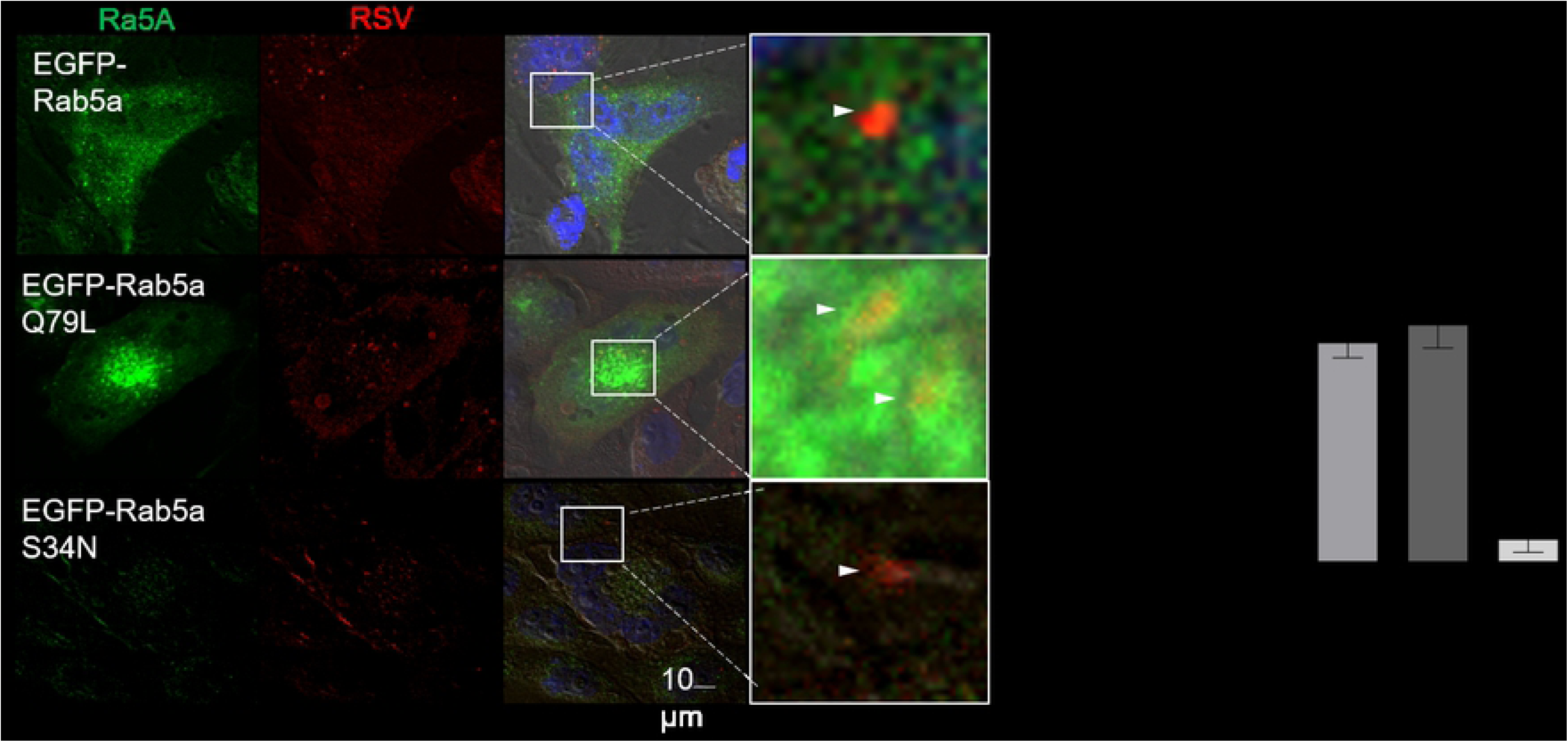
Inactive Rab5a protein decreased RSV replication. A549 cells were transiently transfected with GFP-expressing constructs of Rab5 WT, Rab5 Q79L (C/A), Rab5 S34N (D/N) for 24 h, then infected with RSV for 36 h. (A) The cells were fixed and stained with anti-RSV (red) and anti-DAPI (blue) antibodies. Arrowheads point to RSV and Rab5a co-localization. (B) Viral titers of cell culture homogenates were quantified by plaque assay (plaque forming units, PFU) at 36 h p.i.. \**P* < 0.05; \*\**P* < 0.01; compared with control (RSV, transfected with vector only). Data are shown as mean ± SD of duplicates from at least three independent experiments in duplicate.

### Rab5a depletion exaggerates epithelial antiviral defense to RSV

No mechanism for Rab5a-mediated enhancement of RSV growth has yet been reported. To explore this mechanism, we evaluated the effect of Rab5a signaling on airway epithelial antiviral innate immunity. Type I and Type III IFN play an important role in innate and adaptive antiviral immunity. As indicated earlier, recent studies have implicated that besides IFN-α and IFN-β, IFN-λ is another important IFN that responses to RSV infection in the respiratory epithelia. We, therefore, focused on the potential role for Rab5a in regulating the production of all three IFNs. A549 cells were transfected with siRNAs specific for Rab5a (and siRNA control) for 24 h, and infected with RSV for another 24 h p.i.. Control, mock-infected cells received the same volume of media. Results show that RSV could indeed induce IFN-α (Fig. 4A & 4B), IFN-β (Fig. 4C & 4D) and IFN-λ (Fig. 4E & 4F), when compared with mock infection. Rab5a depletion further increased IFN-λ production significantly (Fig. 4E & 4F), and slightly increased IFN-α and IFN-β production, compared to siCON in RSV-infected cells (Fig. 4A & 4D). The Rab5a S34N mutant also enhanced IFN-λ production (Fig. 4G). These data suggest that during RSV infection Rab5a inhibition increased IFN-λ production.

**FIGURE 4.**
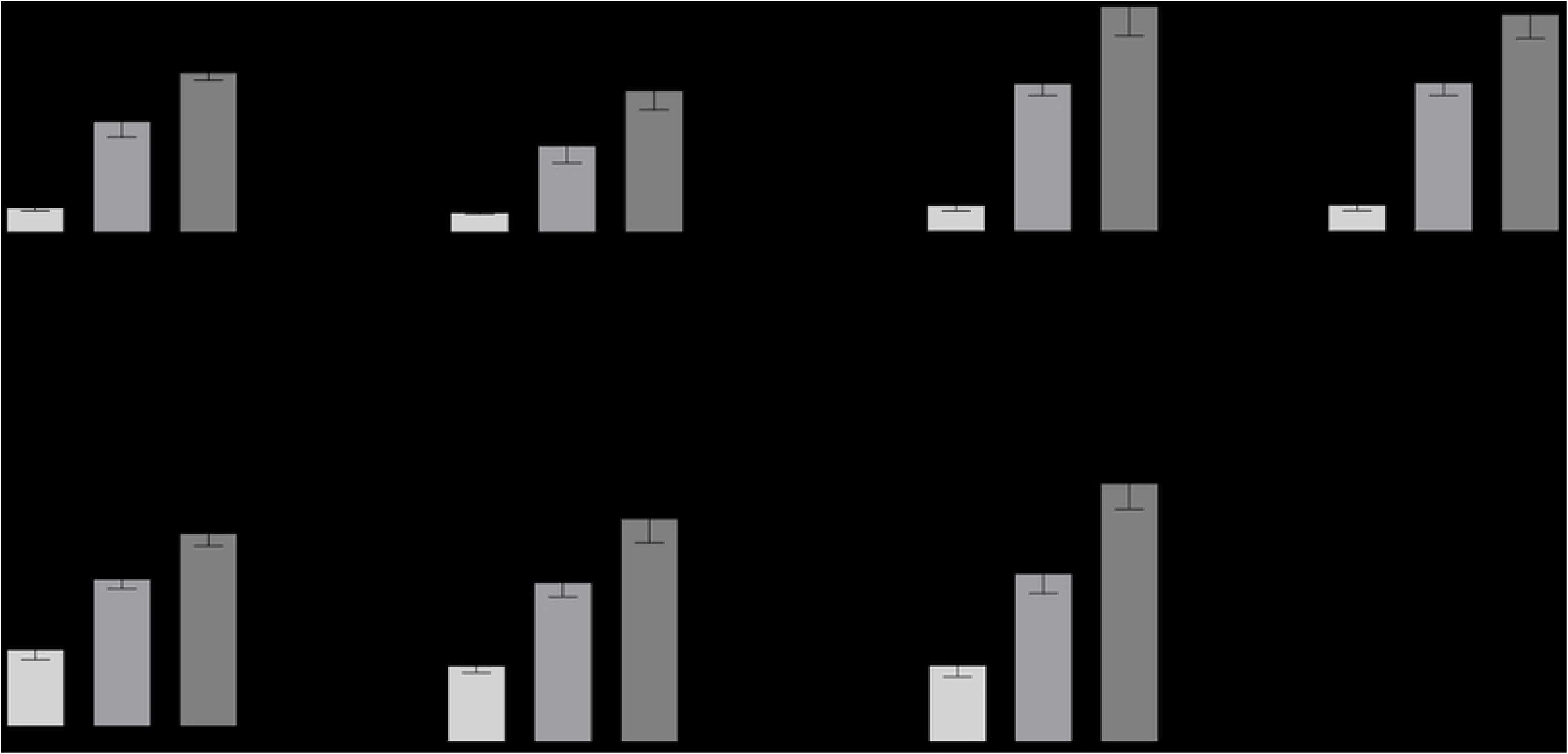
Depletion of Rab5a exaggerates IFN-λ production. Transfection of A549 cells with siRNA control (siCon), Rab5a siRNA, EGFP empty vector (EV) or EGFP S34N, followed by infection with RSV as indicated, have been described in Materials and Methods. All mRNAs were quantified by RT-qPCR, and IFN in the supernatant was quantified by ELISA. (A) IFN-α; (B) IFN-α (IFNA1) mRNA; (C) IFN-β; (D) IFN-β (IFNB1) mRNA; (E) IFN-λ; (F) IFN-λ mRNA. Bars represent the mean ± SD for three independent experiments performed in duplicate. P values were calculated based on Bonferroni of one-way analysis. ***, *p*<0.001, **, *p*<0.01 and *, *p*<0.05 vs. CON and RSV; ^^, *p*<0.01 and ^, *p*<0.05 vs. siCON and siRab5a.

The IFN regulatory factors (IRFs), functioning as transcription factors, play an important role in IFN production. RSV can activate IRF1 in monocytes and lung epithelial cells[42,43], and the IRF1 can interact with the IFN-λ promoter to induce IFN-λ transcription. Thus, to explore the effect of Rab5a on IRF1 expression during RSV infection, we transfected siRab5a and siCON into A549 cells for 24 h, and then infected with RSV for another 24 h. Interestingly, we found that RSV infection increased IRF1 levels, and depletion of Rab5a further increased IRF1 expression during RSV infection (Fig. 5A & 5B). The immunofluorescence staining confirmed these data (Fig. 5C & 5D). Overall, these results suggested that depletion of Rab5a increased IRF1 expression.

**FIGURE 5.**
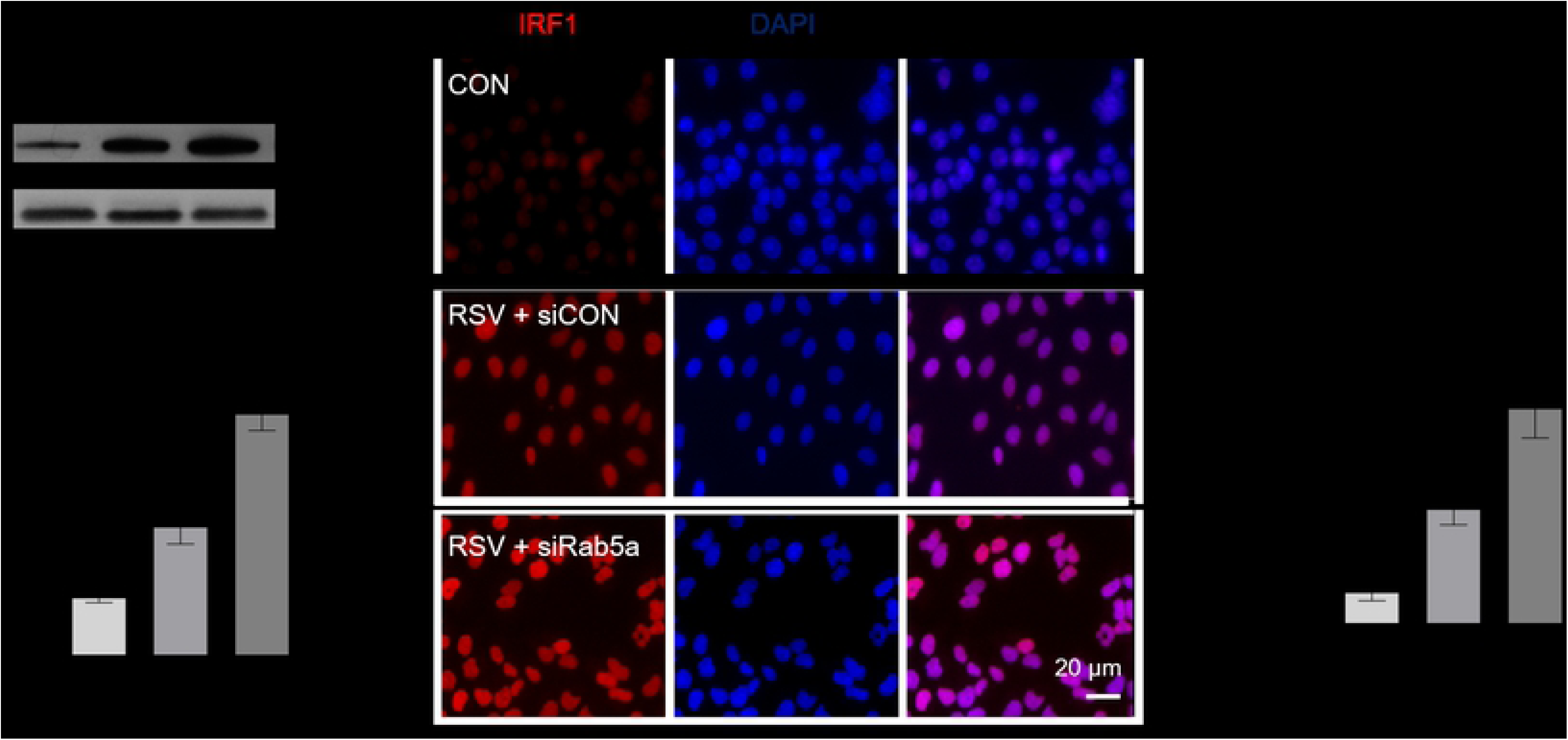
Depletion of Rab5a exaggerates IRF1 production. Transfected with either siRNA control (siCon) or siRNAs targeting IRF1 protein (siIRF1) for 24 h p.i. and infection with RSV have been described in Materials and Methods. (A) IRF1 protein expression, detected by immunoblotting. (B). Semi-quantitative analysis from (A), using Image J. (C) IRF1 protein expression using immunofluorescence assay. (D) Semi-quantitative analysis (mean fluorescence intensity, MFI) from (C), using Image J. Bars represent the mean ± SD for three independent experiments performed in duplicate. P values were calculated based on Bonferroni of one-way analysis. ***, *p*<0.001vs. CON and RSV; ^^^, *p*<0.001 vs. siCON and siRab5a.

To confirm the important role of IRF1 in RSV-induced IFN-λ production in epithelial cells, we treated A549 cells with IRF1-specific siRNA, which significantly suppressed IRF1 protein levels (Fig. 6A). Treatment with this siIRF1 decreased the expression of IFN-λ production in RSV-infected A549 cells, when compared with cells infected with RSV but treated with control siRNA (Fig. 6B). Moreover, siIRF1 eliminated the effect of siRab5a in exaggerating IFN-λ production (Fig. 6C) that we showed earlier (Fig. 4E). Together, these results suggest that: (i) depletion of IRF1 reduces IFN-λ production; (ii) siRab5a exaggerates IFN-λ production via IRF1.

**FIGURE 6.**
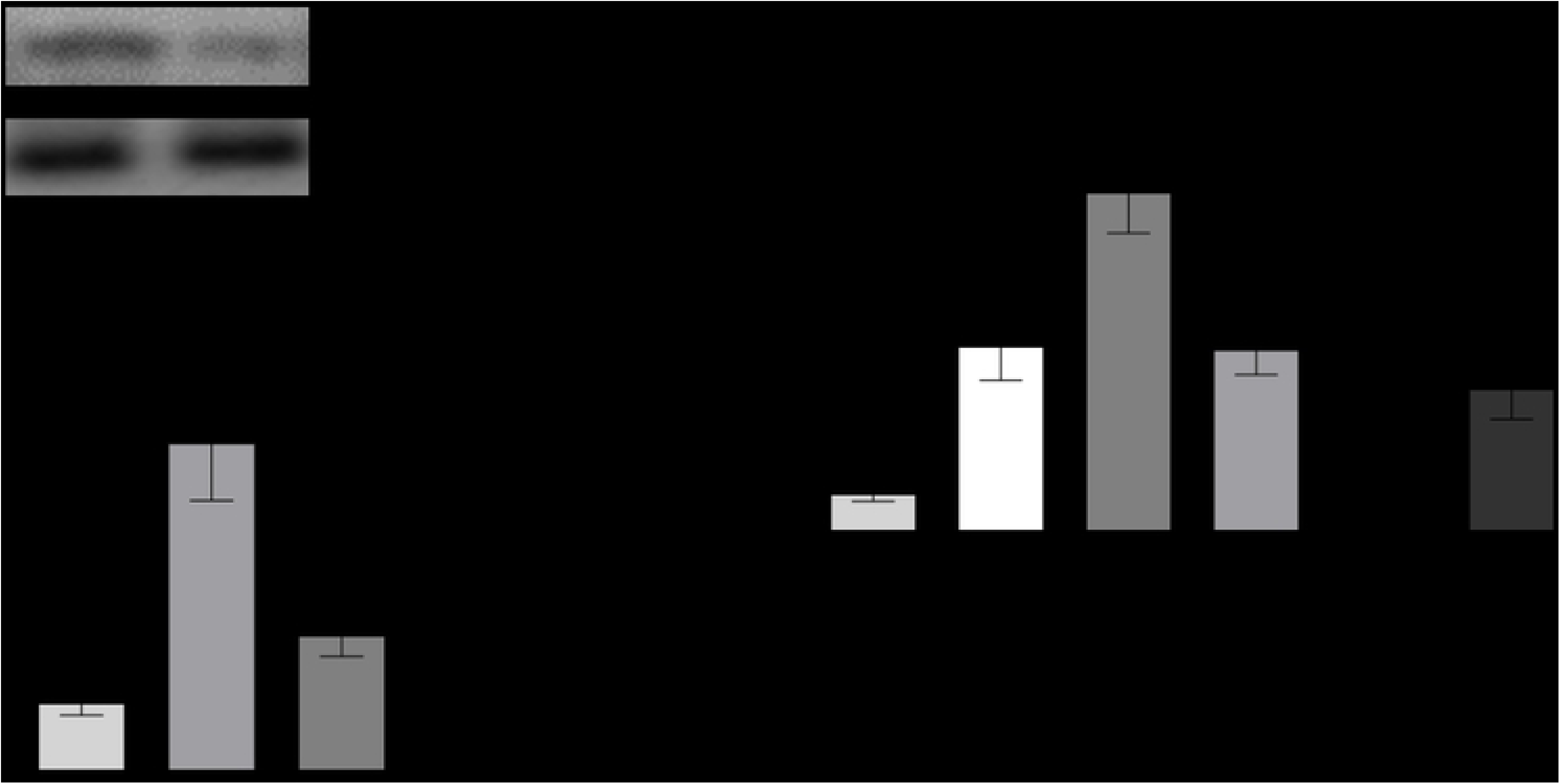
Rab5a mediates IFN-λ production via IRF1. Transfection with siRNA control (siCon) or siRNAs targeting IRF1 protein (siIRF1), and RSV infection, were conducted as described in Materials and Methods. (A) Knockdown efficiency of IRF1 by specific siRNA, determined by immunoblotting. (B) IFN-λ production in supernatants, using ELISA. Bars represent mean ± SD for three independent experiments performed in duplicate. P values were calculated based on Bonferroni of one-way analysis. ***, *p*<0.001, **, *p*<0.01, vs. CON and RSV; ^^, *p*<0.01 vs. siCON and siIRF1 in (B). ***, *p*<0.001, **, *p*<0.01, vs. co-transfected with siRab5a and siIRF1 and transfected with Rab5a or transfected with IRF1 in (C).

### Rab5a depletion leads to an increase of STAT1

IFN-λ binds to its unique receptor complex (IFN-lR1/IL-10R2), which triggers a signaling cascade by activating the downstream JAK-STAT pathway, among which the JAK-STAT1 pathway plays an important role in response to RSV infection. To investigate the effect of Rab5a depletion on JAK-STAT1, A549 cells were transfected with Rab5a-specific siRNA (or control siRNA) for 24 h p.i., then infected with RSV for as before. STAT1 expression was then quantified by immunoblotting. Results (Fig. 7) revealed that RSV infection increased STAT1, and Rab5a depletion increased it further, which lead to an increase in both total STAT1 (Fig. 7AB) and phosphorylated STAT1 species (p-STAT1) (Fig. A,C,D). Moreover, the addition of a JAK1 or STAT1 inhibitor abrogated the ability of siRab5a to suppress RSV infection (Fig. 7D). These results demonstrate that IFN-λ-induced JAK-STAT1 signaling accounts for the effect of IFN-λ on siRab5a-mediated inhibition of RSV.

**FIGURE 7.**
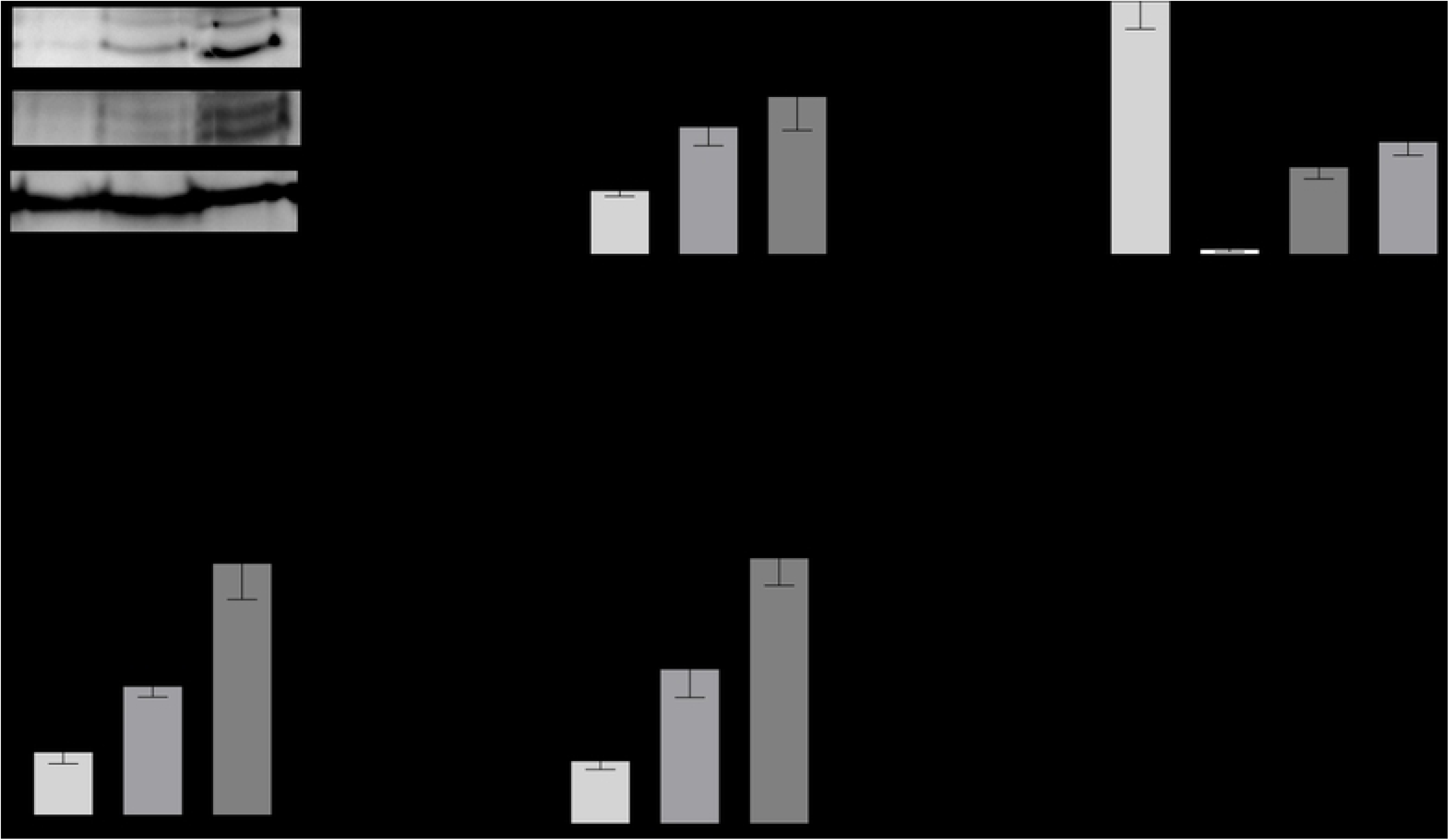
Rab5a depletion activates JAK-STAT1 pathway. siRNA transfection and RSV infection were performed as described in Materials and Methods. (A) Total STAT1 and Phospho-STAT1 (Tyr701) protein expression by immunoblotting. (B) Semi-quantitative analysis of total STAT1 from (A) with Image J. (C, D) Semi-quantitative analysis of phospho-STAT1α or phospho-STAT1β from (A) with Image J. (E) A549 cells were transfected with indicated siRNA, infected with RSV, and treated with JAK1 inhibitor (Baricitinib, 5 nM) or STAT1 inhibitor (Fludarabine, 2.5 μM) for 24 h. Viral titers of cell culture homogenates were assessed by plaque assay as before. Bars represent mean ± SD for three independent experiments performed in duplicate. P values were calculated based on Bonferroni of one-way analysis, ***, *p*<0.001, **, *p*<0.01, vs. CON and RSV. P values were calculated based on unpaired Student’s t-test, ^^, *p*<0.01 vs. siCON and siRab5a in (B, D). ^^^, *p*<0.001 vs. siRab5a and siRab5a with Baricitinib or siRab5a with Fludarabine in (E).

### Rab5a depletion amplifies RIG-I and Mx1 expression, in part via the JAK-STAT1 pathway

Previous studies found that RSV infection induces RIG-I and Mx1 production[44,45]. RIG-I and Mx1 are downstream genes of the JAK-STAT1-dependent IFN response pathway. We first confirmed this induction, and showed that depletion of Rab5a indeed increased the expression of these two genes (Fig.8A & 8C). Inhibition of JAK and STAT1 by specific inhibitors, Baricitinb and Fludarabine respectively, partially rescued this increase of RIG-I and Mx1 (Fig. 7B & 7D), suggesting that the induction of RIG-I and Mx1 by Rab5a occurs via the JAK-STAT pathway.

## Discussion

Results presented here support an important role of Rab5a in RSV replication. RSV infection increased the amount of Rab5a in airway epithelial cells. The biological effect of this increased Rab5a is to suppress host anti-viral immunity (Fig. 9A). Knockdown of Rab5a gene expression or its inhibition by dominant negative mutants in A549 cells results in protection against RSV infection through the activation of IRF1-dependent, IFN-λ mediated anti-viral pathway (Fig. 9B). Indeed, the presence of Rab5a leads to higher infection and replication of the virus. This study is the first to show that in airway epithelial cells: (i) RSV infection alters the amount of Rab5a, (ii) Rab5a enhances RSV replication, (iii) the mechanism of this effect of Rab5a involves regulation of IRF1, IFN-λ and STAT1, and (iv) Rab5a may thus attenuate inflammation of the epithelia during RSV infection.

**FIGURE 8.**
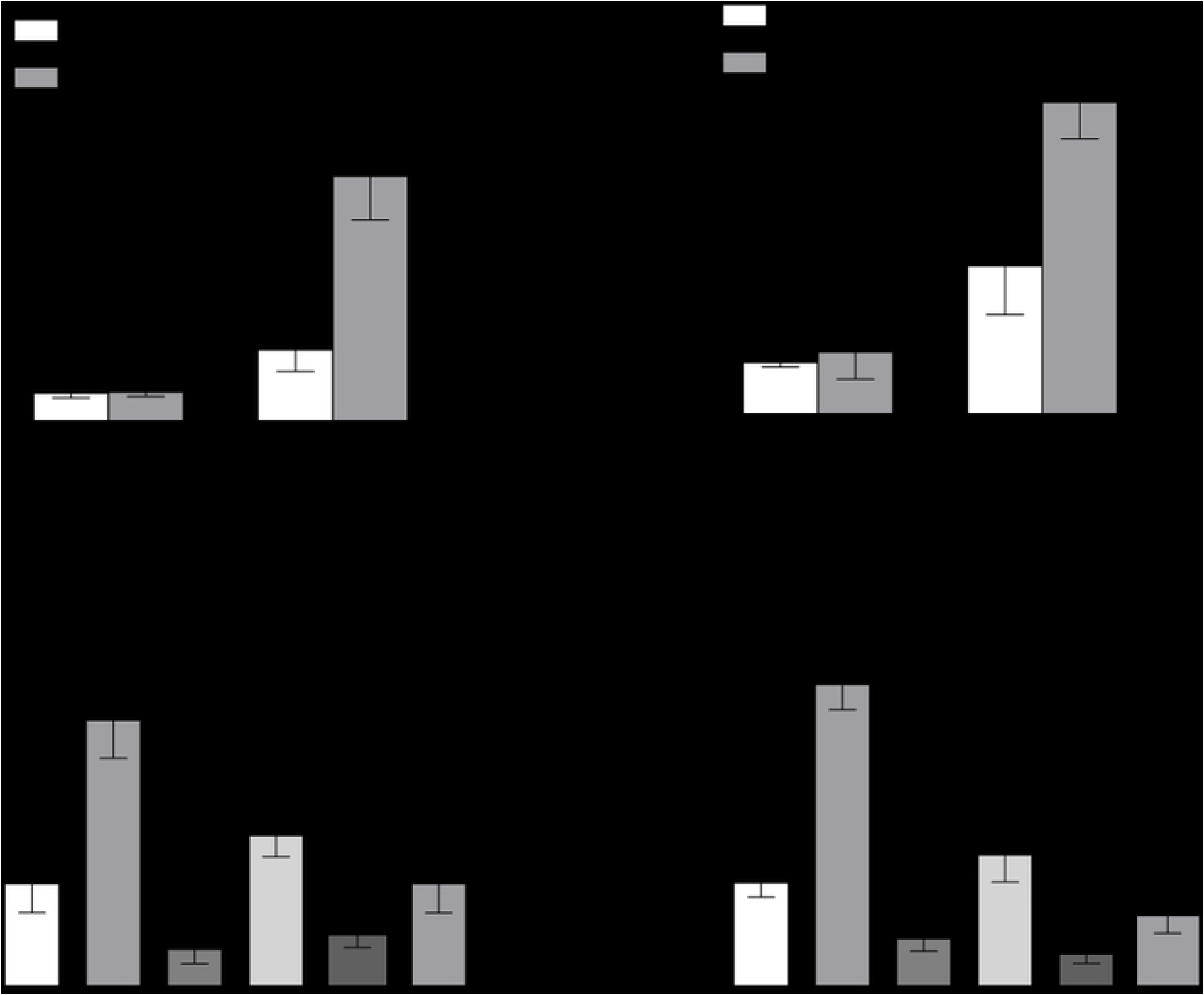
Rab5a depletion increases RIG-I and Mx1 mRNA expression. Transfection of A549 cells and RSV infection were performed as before. (A, C) RIG-I and Mx1 mRNA expression, measured by RT-qPCR. (B, D) Where indicated JAK1 inhibitor (Baricitinib, 5 nM) or STAT1 inhibitor (Fludarabine, 2.5 μM) were used for 24 h; RIG-I and Mx1 mRNA were quantified by RT-qPCR. Bars represent mean ± SD for three independent experiments performed in duplicate. P values were calculated based on Bonferroni of one-way analysis, ***, *p*<0.001, **, *p*<0.01, vs. CON and RSV. P values were calculated based on unpaired Student’s t-test, ^^, *p*<0.01 vs. siCON and siRab5a. ###, *p*<0.001, ##, *p*<0.01 vs. siRab5a and siRab5a with Baricitinib or siRab5a with Fludarabine.

**FIGURE 9.**
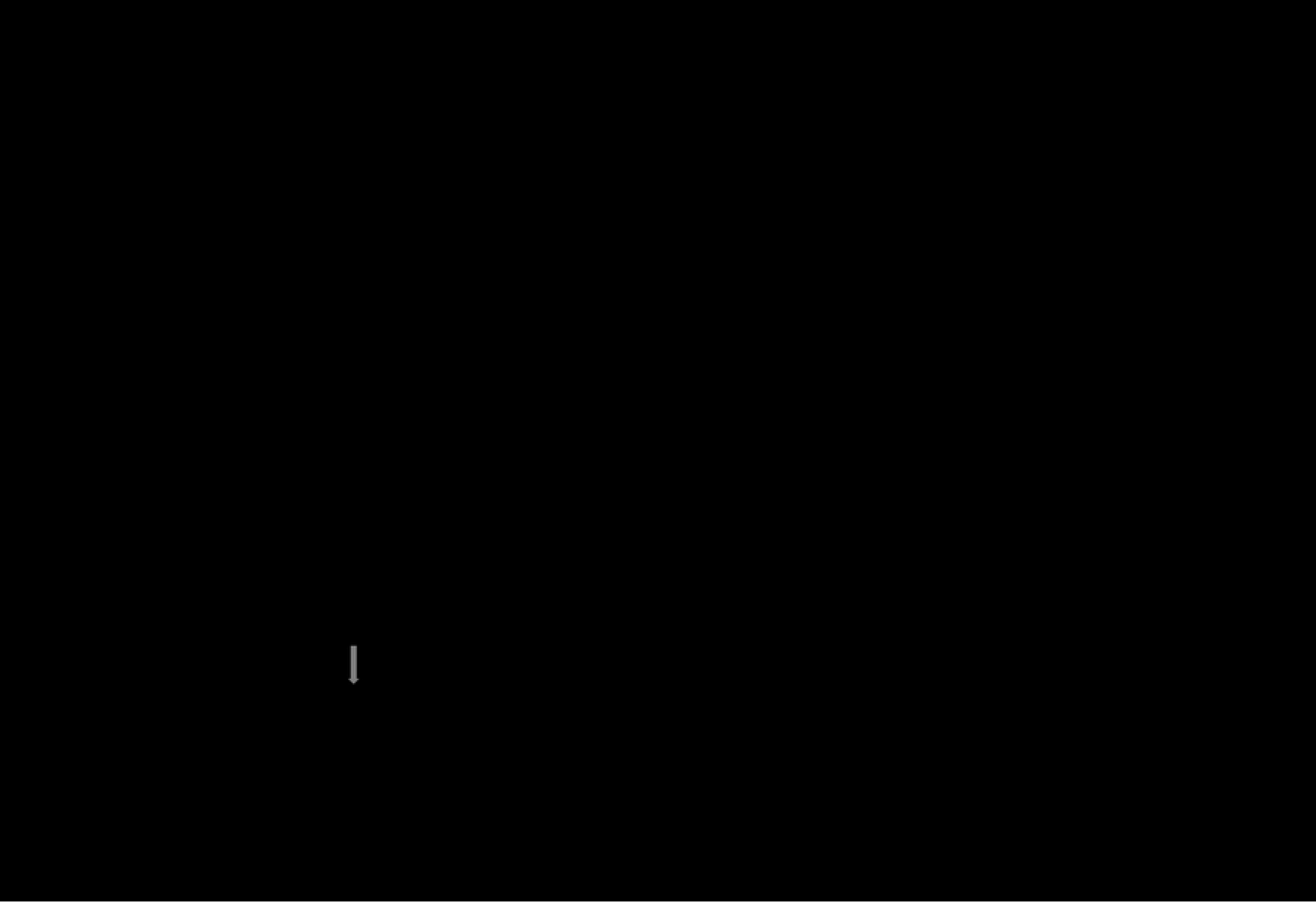
Overview. (A) RSV activates Rab5a that inactivates IRF1. IRF1 inactivation inhibits IFN-λ production, which decreases JAK/STAT1 pathway activation, resulting in suppression of host defense and increased viral replication. (B) Upon Rab5a depletion (by siRNA), IRF1-induced IFN-λ is increased, which results in decreased viral titers.

While previous studies have demonstrated an effect of Rab5a in diverse virus replication [18,21,46], none have shown an alteration of Rab5a protein levels in RSV infection. Krzyzaniak et al found that inhibiting the activation of Rab5a attenuates viral titer in RSV infection of HeLa cells [8]. Another study, mentioned earlier, found that depletion of Rab5a resulted in decreased RSV replication, and viral binding was not affected when suppressed action of Rab5a [9]. We confirmed and extended these studies and found that there is no significant difference of viral binding between Rab5a-depleted and untreated cells (Fig. S3). Together, these results establish that depletion of Rab5a has no effect on RSV binding to airway epithelial cells.

Rab5a is also required for macrosome formation. Previous studies suggested that RSV entry into host cell takes place via actin-related micropinocytosis [8]. In the very first step of RSV infection, the virions need to engage cellular receptors to trigger entry into the cell. Studies have suggested a plethora of candidate cellular receptors for RSV entry, for example, CX3CR [47–49], EGFR [50,51], TLR4 [52,53], ICAM-1 [54], nucleolin [55,56], and HSPGs [57]. Among these, EGFR plays a particularly important role in RSV infection and inflammation [58], and the mechanism is related to IRF1-dependent IFN-λ [35]. Multiple studies have shown that EGFR closely interacts with Rab5a. For example, Rab5a is very important for EGFR trafficking, and endogenous EGFRs can partially co-localize with endogenous Rab5 [59]. Stahl [60] reported that EGFR stimulates the activation of Rab5a, and also enhances the translocation of Rab5a. Meanwhile, suppression of Rab5a hampered the degradation of EGFR and its internalization. Furthermore, Rab5a overexpression facilitated cell proliferation through the EGFR signal pathway [61]. In our studies, RSV infection activated not only Rab5a (Fig. 2), but also EGFR (unpublished data). Depletion of Rab5a decreased the activation of EGFR, and inhibition of EGFR by Gefitinib also suppressed Rab5a protein level; finally, both Rab5a depletion and Gefitinib treatment decrease RSV infection via IRF1-dependented IFN-λ production [35]. Altogether, our current studies predict that Rab5a may promote viral infection through EGFR, further investigation is needed to determine how exactly Rab5a and EGFR influence each other during RSV infection.

The nonstructural proteins (NS1 and NS2) of RSV suppress host innate and adaptive immune responses against the virus. Both proteins, individually and in combination, inhibit type I IFN pathway [62], and the NS1 protein also suppresses IFN-λ production in airway epithelial cells during RSV infection [37]. In our study, depletion of Rab5a amplified IFN-λ production in RSV infection, moreover, overexpression of Rab5a increased NS1 mRNA and protein level (data not shown). Therefore, our data predicted that Rab5a also may promote RSV NS1 production to anti-epithelial antiviral defenses. We need further investigate the correlation between Rab5a and RSV NS1.

Lastly, we have documented that inhibition of Rab5a needed IRF1and IFN-λ to restrain RSV infection, and the JAK-STAT1 pathway was implicated in this effect. In agreement with a previous study [35], we also found that exogenously added recombinant IFN-λ could inhibit RSV infection, which demonstrate an important antiviral role of this pathway, defending the airway epithelial cells against RSV. However, the mechanism by which Rab5a suppresses IRF1still needs to be elucidated. Another study found that ERK inhibition could increase RSV-induced IFN-λ production [35]. Therefore, we need further investigate whether ERK signaling may act as a critical link in the mechanism connecting IRF1 and Rab5a.

In conclusion, although limited in scope, our studies have demonstrated that Rab5a plays an important role in RSV infection. We have also discovered a new mechanism in which RSV uses Rab5a to suppress epithelial antiviral immunity, such that silencing or inactivating Rab5a results in reduced viral infection. This is a new insight on the role of the cellular factor Rab5a in RSV infection, which can be explored as a therapeutic and druggable target against RSV infection.

## Materials and Methods

### Research subjects

Twenty-six children with RSV infection, hospitalized from November 2016 to January 2017 in the Department of Respiratory Medicine, Children’s Hospital, Chongqing Medical University, were enrolled in this study. Nasopharyngeal aspirates (NPAs) were prospectively collected from all subjects within the first day after hospital admission. Immunofluorescence assays were performed to detect the presence of RSV, adenovirus, influenza A and B virus, and parainfluenza virus 1, 2, and 3 in the NPAs. Infants that carried viruses other than RSV were excluded from the study. Those considered positive for bacterial infection on the basis of published criteria [63] were excluded as well. The control group was selected from infants with no evidence of infection, and underwent surgical therapy only to clear secretions from the airway.

### Nasopharyngeal aspirate analysis

NPAs were collected and analyzed as previously described [63]. In brief, they were collected gently and Mxed uniformly. To 0.5 mL of the aspirate, transferred to a new tube, 2 mL of 0.1% dithiothreitol was added. The Mxture was vortexed three times, 15 seconds each, and rocked on a bench rocker for 15 min. The suspension was collected and subsequently centrifuged at 306 g for 10 min. Cell-free supernatants were collected and aliquots stored at –80°C. The cell pellet was used for total RNA and protein extraction.

### Reagents, antibodies and plasmids

The primary antibodies used in this study include a goat polyclonal antibody to RSV, purchased from Millipore, and rabbit Rab (Rab1a, Rab2a, Rab4a, Rab5a, Rab6a, Rab7a, Rab8a, Rab9a and Rab11a), STAT1, phospho-STAT1 (Tyr701), IRF1, and mouse GAPDH monoclonal antibodies, purchased from Cell Signaling Technology (CST), USA. DAPI was purchased from Sigma-Aldrich, USA. The secondary antibodies used were Alexa Fluor 568/488-conjugated duck anti-goat IgG or anti-rabbit IgG from Biyuntian, Beijing, China, and horseradish peroxidase (HRP)-conjugated goat anti-rabbit IgG or anti-mouse IgG from CST, USA. Expression plasmids encoding EGFP-tagged Rab5a and its mutants were purchased from Addgene, USA. The Janus kinase 1 (JAK1) inhibitor, Baricitinib, was purchased from MedChemExpress, Shanghai, China, and the STAT1 inhibitor, Fludarabine, from Selleckchem.

### Cell culture, virus and infection

Human alveolar carcinoma type II-like epithelial cell line A549 (ATCC CCL-185) and human laryngeal cancer epithelial cell line HEp-2 (ATCC CCL-23) were cultured in Dulbecco’s Modified Eagle Medium (DMEM) supplemented with 10% fetal bovine. RSV A2 strain was obtained from ATCC. For all experiments, RSV was grown in HEp-2 cells with 5% FBS and purified by density gradient as previously described [64]. In all experiments where RSV infection was performed, a multiplicity of infection (MOI) of 0.8 was used.

### Virus titration

At indicated times, the infected cell media supernatants were collected, and the cells were scraped into the cell culture medium and vortexed three times with glass beads, followed by centrifugation at 1000 rpm for 5 min. RSV titration was performed by plaque assay on HEp-2 cells.

### siRNA transfection

The Rab (Rab1a, Rab4a, Rab5a, Rab6a, Rab7a, Rab8a, Rab9a and Rab11a) siRNA sequences were from reference [65], and were as follows. The Rab1a siRNAs: 5’CAGCAUGAAUCCCGAAUAU; 5’GUAGAACAGUCUUUCAUGA; 5’ GUAGAACAGUCUUUCAUGA; 5’UGAGAAGUCCAAUGUUAAA; Rab4a siRNAs :5’GAAAGAAUGGGCUCAGGUA; 5’GUUAACAGAUGCCCGAAUG; 5’UUAGAAGCCUCCAGAUUUG; 5’UACAAUGCGCUUACUAAUU; Rab5a siRNAs: 5’GCAAGCAAGUCCUAACAUU; 5’GGAAGAGGAGUAGACCUUA; 5’AGGAAUCAGUGUUGUAGUA; 5’GAAGAGGAGUAGACCUUAC; Rab6a siRNA: 5’GAGAAGAUAUGAUUGACAU; 5’GAGCAACCAGUCAGUGAAG; 5’AAGCAGAGAAGAUAUGAUU; 5’CCAAAGAGCUGAAUGUUAU; Rab7a siRNA: 5’GGGAGUUCUGGAGUCGGGAA; 5’CCACAAUAGGAGCUGACUU3’. Rab8a siRNAs: 5’GAAUUAAACUGCAGAUAUG; 5’GAACAAGUGUGAUGUGAAU; 5’GAACUGGAUUCGCAACAUU; 5’GAAGACCUGUGUCCUGUUC; Rab9a siRNA duplex: 5’CGGCAGGTGTCTACAGAAG; Rab11A siRNAs: 5’GGAGUAGAGUUUGCAACAA; 5’GUAGGUGCCUUAUUGGUUU; 5’GCAACAAUGUGGUUCCUAU; 5’CAAGAGCGAUAUCGAGCUA. These and the IRF1 siRNA (duplex UCCCAAGACGUGGAAGGCCAACUUU) were purchased from Qiagen. The siRNA against human Rab2a was purchase from Sigma Aldrich. Cell lysates were separated by SDS-PAGE and subjected to immunoblotting with rabbit anti-Rabs, rabbit anti-IRF1, or mouse anti-GAPDH (as a loading control) antibodies.

### Immunoblotting

Total cell protein (50 μg) was resolved on 5-15% Bis-tris SDS-PAGE, and transferred onto PVDF membranes. The membranes were incubated with primary antibodies against Rab proteins, IRF1, GAPDH, or LaminB1, followed by incubation with appropriate secondary antibodies. The protein bands were visualized with a chemiluminescence kit (Bio-Rad).

### Quantitative (q) RT-PCR

A549 cells, transfected with plasmids, and infected with RSV as and where mentioned, were harvested, and intracellular RNA was purified at indicated times post-transfection (BioTake). Viral RNA was isolated from the Mxture of cells and media supernatants. The RNA (1 μg) was used for first-strand cDNA synthesis and the cDNA was amplified using the VeriQuest Fast SYBR Green qPCR kit (invtrogen) with primers as follows: Rab5a (forward primer 5’ CAAGAACGATACCATAGCCTAGCAC3’; reverse primer 5’ CTTGCCTCTGAAGTTCTTTAACCC 3’); IFNA1 (forward primer 5’ GTGAGGAAATACTTCCAAAGAATCAC3’; reverse primer 5’ TCTCATGATTTCTGCTCTGACAA 3’); IFNB1 (forward primer 5’ CAGCAATTTTCAGTGTCAGAAGC3’; reverse primer 5’ TCATCCTGTCCTTGAGGCAGT 3’); IFN-λ(forward primer 5’ CGCCTTGGAAGAGTCACTCA3’; reverse primer 5’ GAAGCCTCAGGTCCCAATTC3’).

RSV copy numbers were quantified with TaqMan RT-PCR as previously described [66]. The PCR cycle conditions were as follows: 50°C for 2 min, 95°C for 10 min, 40 cycle at 95°C for 15 s and 60°C for 30 s. The fold change was obtained using 2-∆∆Ct method using GAPDH as a calibrator.

### Transfection and transient expression

All plasmid transfections were performed with a commercial transfection kit (Thermo Fisher Scientific, USA). Cells were seeded on 12 mm coverslips in 24 wells for imaging or in 6-well plates for qRT-PCR or immunoblotting analyses. Experiments were performed at indicated times after transfection.

### Indirect immunofluorescence assays

RSV-infected A549 cells were fixed with 4% paraformaldehyde for 30 min at room temperature and permeabilized with 0.2% Triton X-100 (sigma) for an additional 20 min. After washing and blocking with 2% BSA for 1 h, the cells were incubated with anti-IRF1 or anti-RSV antibody at 4 ^0^C overnight. The cells were then washed three times, followed by incubation with Alexa Fluor 568-conjugated secondary antibodies (Molecular Probes) for 1 h at room temperature. Finally, the cells were visualized and photographed using a Nikon laser confocal microscope. in which the images were acquired with confocal Z-section series and were subsequently analyzed with NIS-Elements BR software, version 4.11.

### Syncytia quantification

Quantification of the RSV syncytia was performed as described by Buchholz et al (2016)[67], with minor modifications. A549 cells were grown in three coverslips in 24 wells per group, and infected with RSV at MOI 0.8 where mentioned, then fixed at 24 h post-infection (p.i.), and stained with DAPI and for RSV. The three coverslips were imaged and NIS software was used to draw the area of the syncytia. Image J (NIH) was used to confirm the syncytial area. Syncytia were counted when they were RSV-positive.

### Measurement of IFN

Human-specific enzyme-linked immunosorbent assay (ELISA) kits were used to measure IFN-α, IFN-β and IFN-λ levels in culture supernatants (BD, USA).

### Statistical analysis

Data analysis were performed using SPSS 19.0 software. One-way analysis of variance (ANOVA) was used to detect the significance of the difference among groups. Unpaired Student’s t-test was used to detect the significance between two groups. *P* value of < 0.05 was considered significant.

### Ethics Statement

Use of NPA samples of infants were approved by the Ethics Committee of the Children’s Hospital, Chongqing Medical University (permit number 2015–77). The parents or legal guardians offered written informed consent to participate in the study before the infants were enrolled. All procedures were performed in accordance with the approved guidelines, and obeyed the principles of the Declaration of Helsinki.

## ACKNOWLEDGMENTS

This research is supported by grants from the National Natural Science Foundation of China 81670011, 91642107.

